# The tetrapeptide sequence of IL-1β regulates its recruitment and activation by inflammatory caspases

**DOI:** 10.1101/2023.02.16.528859

**Authors:** Patrick M. Exconde, Claudia Hernandez-Chavez, Mark B. Bray, Jan L. Lopez, Tamanna Srivastava, Marisa S. Egan, Jenna Zhang, Sunny Shin, Bohdana M. Discher, Cornelius Y. Taabazuing

**Author notes:** Patrick M. Exconde and Claudia Hernandez-Chavez contributed equally to this work. Correspondence to Cornelius Y. Taabazuing.

## Abstract

The mammalian innate immune system uses germline-encoded cytosolic pattern-recognition receptors (PRRs) to detect intracellular danger signals. At least six of these PRRs are known to form multiprotein complexes called inflammasomes which activate cysteine proteases known as caspases. Canonical inflammasomes recruit and activate caspase-1 (CASP1), which in turn cleaves and activates inflammatory cytokines such as IL-1β and IL-18, as well as the pore forming protein, gasdermin D (GSDMD), to induce pyroptotic cell death. In contrast, non-canonical inflammasomes, caspases-4/-5 (CASP4/5) in humans and caspase-11 (CASP11) in mice, are activated by intracellular LPS to cleave GSDMD, but their role in direct processing of inflammatory cytokines has not been established. Here we show that active CASP4/5 directly cleave IL-18 to generate the active species. Surprisingly, we also discovered that CASP4/5/11 cleave IL-1β at D27 to generate a 27 kDa fragment that is predicted to be inactive and cannot signal to the IL-1 receptor. Mechanistically, we discovered that the sequence identity of the P4-P1 tetrapeptide sequence adjacent to the caspase cleavage site (D116) regulates the recruitment and processing of IL-1β by inflammatory caspases to generate the bioactive species. Thus, we have identified new substrates of the non-canonical inflammasomes and reveal key mechanistic details regulating inflammation.

## Introduction

The mammalian innate immune system uses germline-encoded pattern recognition receptors (PRRs) to detect pathogen-associated molecular patterns (PAMPs) or damage-associated molecular patterns (DAMPs)^1,2^. Upon detecting PAMPS or DAMPs, some of these PRRs rapidly assemble into multi-protein complexes termed inflammasomes^1,2^. Canonical inflammasome assembly typically involves recruitment of the adapter protein ASC and then pro-caspase-1 (pro-CASP1) to the PRR, which leads to oligomerization and auto-proteolytic maturation of pro-CASP1 into the active species. Active CASP1 then processes the inflammatory cytokines IL-1β and IL-18 into their bioactive forms, as well as the pore forming protein, gasdermin D (GSDMD)^3–5^. GSDMD processing liberates the N-terminus from the inhibitory C-terminal fragment, allowing the N-terminus to oligomerize and form pores in the plasma membrane to facilitate the release of cytokines and induce pyroptotic cell death^6–8^.

During canonical inflammasome activation, pro-CASP1 undergoes auto-proteolysis at multiple sites to generate distinct species (**Fig. 1A**)^9–12^. It is thought that IDL processing confers full protease activity to the inflammatory caspases^9,12–14^. In support of this idea, recent studies suggest that the CASP1 p33/10 species is the active species in cells and the p20/10 rapidly loses protease activity^9^. However, ASC, which facilitates full autoproteolysis of pro-CASP1 into the p20/10 species, is required for IL-1β processing for reasons that are not clear^10,11^. Current structural studies also suggest that the CASP1 p20/10 is the species that binds GSDMD^15,16^. Of note, CASP1 p20/10 was demonstrated to bind GSDMD through exosite interactions, but how CASP1 recognizes other substrates such as IL-1β and IL-18 remains an open question.

**Figure 1.**
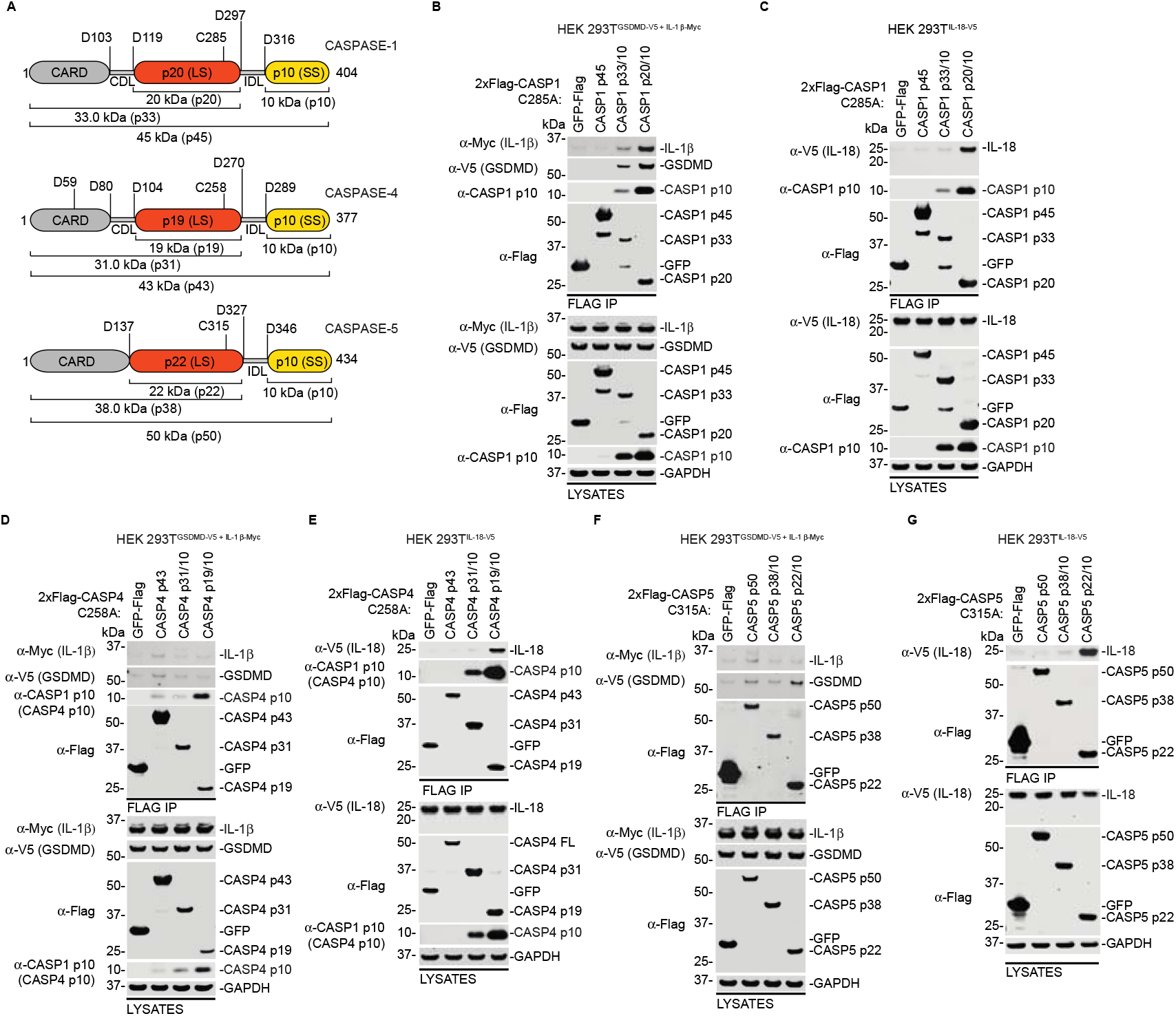
The p20/10 species of human inflammatory caspases are the species that bind strongly to inflammatory substrates. (**A**) Schematic of human CASP1 (top), CASP4 (middle) and CASP5 (bottom) depicting the catalytic cysteines and autoproteolytic sites that give rise to the distinct caspase species. (**B**,**D**,**F**) HEK 293T cells stably expressing C-terminally V5-tagged GSDMD (GSDMD-V5) and Myc-tagged IL-1β (IL-1β-Myc) were transiently transfected with the indicated catalytically inactive caspase constructs. After 48 h, the cells were harvested and subjected to anti-FLAG IP followed by immunoblot analysis. GFP control was C-terminally FLAG tagged (GFP-FLAG) and the catalytically dead caspases all harbored an N-terminal 2xFLAG tag. (**C**,**E**,**G**) HEK 293T cells stably expressing C-terminally V5-tagged IL-18 (IL-18-V5) were transiently transfected with the indicated constructs as in B,D, and F for 48 h before the cells were harvested and subjected to anti-FLAG IP followed by immunoblotting. Data are representative of three or more independent experiments.

In contrast to the canonical inflammasome pathway, non-canonical inflammasome activation involves direct detection of cytosolic lipopolysaccharide (LPS) from gram-negative bacteria by caspases-4/5 (CASP4/5) in humans and caspase-11 (CASP11) in mice^17–20^. Binding of LPS induces oligomerization and autoproteolytic maturation of the non-canonical inflammasomes (CASP4/5/11) to generate the active proteases^13,14,21^. Active CASP4/5/11 then cleave GSDMD to induce pyroptotic cell death. Formation of GSDMD pores during non-canonical inflammasome activation results in K^+^ efflux, which activates CASP1 to induce IL-1β and IL-18 processing via the canonical NLRP3 inflammasome ^22,23^. Interestingly, both CASP4 and CASP11 have been implicated in IL-18 processing, but as the canonical pathway is activated downstream, whether this is a direct processing event in cells or if CASP5, which is thought to be functionally similar to CASP4, can analogously process IL-18 remains to be defined^24–27^. Furthermore, like CASP1, autoproteolyzed CASP4/11 were also reported to utilized exosite mediated interactions for GSDMD binding, but if and how the non-canonical inflammasomes interact with other putative substrates such as IL-18 is unknown. The contributions of the different inflammatory caspase species and mechanisms employed for substrate recognition and processing need to be clearly defined as this has important ramifications for innate immune regulation.

Here, we used a combination of biochemistry and chemical biology to uncover the substrates cleaved by specific caspase species and to test the role of non-canonical inflammasomes in cytokine maturation. We discovered that distinct inflammatory caspase species interact with specific substrates with varying affinities, which likely regulates substrate processing. Interestingly, we discovered that both CASP4/5 cleave IL-18 directly, but CASP11 is unable to cleave IL-18. Surprisingly, we also discovered that CASP4/5/11 cleave IL-1β to generate a 27 kDa fragment that was previously reported to abrogate signaling to the IL-1 receptor^28^. Importantly, we demonstrate that the inflammatory caspases recognize the P4–P1 tetrapeptide sequence adjacent to the IL-1β processing site (D116) to facilitate binding and processing. Altogether, we identified new direct substrates of the non-canonical inflammasome pathway and uncovered the molecular regulation governing cytokine processing by the inflammatory caspases.

## RESULTS

### The p20/10 species of human inflammatory caspases are the species that bind strongly to inflammatory substrates

The inflammatory caspases are comprised of a CARD domain, followed by a 20 kDa large catalytic subunit (LS), and a 10 kDa small subunit (SS) (**Fig. 1A**). During activation,autoproteolytic maturation first occurs at the interdomain linker (IDL) which joins the LS and SS to generate the active p33/10 species for CASP1, p31/10 species for CASP4, and a p38/10 species for CASP5 (**Fig. 1A**)^9,12^. Because IDL cleavage generates a CARD-LS that is ∼35 kDa when averaged, we will henceforth refer to the p35/10 when referencing them collectively for conceptual ease. Next, autoproteolysis at the CARD-domain linker (CDL), which joins the CARD-domain and LS, generates the fully matured p20/10 species for CASP1, p19/10 species for CASP4, and p22/10 species for CASP5 (**Fig. 1A**). We will hereafter refer to them as the p20/10 species when discussing them collectively. As these distinct species may have different substrate specificities, we first wanted to determine at what point during activation does the inflammasome interact with specific substrates. Expression of the large and small subunits of inflammatory caspases as separate polypeptides results in formation of the active species^16^. We first generated plasmids that allowed us to express the different subunits of CASP1/4/5 as separate polypeptides using a single plasmid. This granted us the ability to express the different active species of each inflammatory caspase in the absence of any stimulating ligand for functional interrogation. We transiently expressed the catalytically inactive (cysteine to alanine mutants) of the different CASP1/4/5 species harboring a 2xFLAG N-terminal tag in HEK 293T cells ectopically expressing GSDMD-V5 and IL-1β-Myc, or HEK 293T cells expressing IL-18-V5 (**Fig. 1B-G**). The lysates were collected and subjected to anti-FLAG immunoprecipitation. As expected, the CASP1 p20/p10 species bound GSDMD, IL-β (**Fig. 1B**), and IL-18 (**Fig. 1C**). GSDMD and IL-1β were bound weakly to CASP1 p33/10 but we could not detect any binding of IL-18 to CASP1 p33/10 (**Fig. 1B,C**).

We then probed for binding of the human non-canonical caspases to the inflammatory substrates (GSDMD, IL-1β, and IL-18). We could not detect binding of GSDMD and IL-1β to any of the CASP4 species tested in this assay (**Fig. 1D**). However, we detected binding of IL-18 to CASP4 p19/10 (**Fig. 1E**). During this work, we noticed that the CASP1 p12/10 antibody (Abcam) had significant cross reactivity with the CASP4 p10 subunit. We therefore used this antibody to check the expression of CASP4 p10 as no commercially available antibody exists that detects the p10. Unfortunately, none of the CASP5 antibodies we tried detected the p10 subunit. Notably, CASP4 p10, which was expressed separately, was pulled down by the p19 subunit, suggesting that a functional enzyme complex was formed, but the complex did not interact strongly with GSDMD and IL-1β. Similarly, IL-18 was bound to CASP5 p22/10 and we detected modest binding of GSDMD, but not IL-1β to CASP5 p22/10 (**Fig. 1F,G**). It is worth noting that no processing of GSDMD, IL-1β, or IL-18 was observed with expression of the catalytically inactive caspase species. Taken together, this data suggests that the auto proteolyzed p20/10 species of CASP4/5 binds strongly to IL-18, but only form weak interactions with GSDMD and IL-1β.

### CASP4/5 cleave IL-1β and IL-18

We next sought to investigate the functional impact of the distinct caspase species on inflammatory substrate processing. We hypothesized that when the distinct species are present at the same levels in cells, the p20/10 species would be the most active species given that they exhibited the strongest binding to the inflammatory substrates. We thus expressed the catalytically competent p35/10 and p20/10 constructs of CASP1, −4, and −5 as separate polypeptides in the HEK 293T cells stably expressing the inflammatory substrates described above. To prevent processing of the p35 constructs, we made the corresponding CDL mutations for CASP1 (D103A, D119A), CASP4 (D59A, D80A, D104A), and CASP5 (D137A) to generate constitutive p35/10 enzymes. Transient transfection of the p35/10 or p20/10 species in HEK 293T cells expressing GSDMD and IL-1β resulted in significant LDH release for all species (**Fig. 2A**) except for CASP5 p20/10, which was expressed at lower levels (**Fig. 2B**). Notably, the CASP1 and CASP4 p35/10 and p20/10 species were expressed at relatively comparable levels, but we consistently saw less expression of CASP5 p22/10 species, suggesting CASP5 p22/10 may be unstable when expressed. In agreement with the p20/10 species being the species that bind strongly to the inflammatory substrates, we observed slightly higher LDH release for these species compared to the p35/10 species (**Fig. 2A**). Intriguingly, all the active species seemed to induce some apoptosis as evidenced by PARP cleavage, but only the CASP1 p33/10 and p20/10 species robustly processed IL-1β and GSDMD (**Fig. 2B**). Both the CASP4 p31/10 and p19/10 species had some slight GSDMD processing, but we could not detect GSDMD processing for CASP5 (**Fig. 2B**). Surprisingly, CASP4 also cleaved IL-1β to generate the active 17 kDa fragment, but this processing event was minor compared to CASP1 (**Fig. 2B**). Unexpectedly, we also observed a 27 kDa cleavage product of IL-1β in CASP4 and CASP5 transfected cells (**Fig. 2B**). We note that CASP5 p22/10 was expressed less in this assay, and although we could not detect the p10 subunit using an antibody, we could confirm expression based on functional processing of substrates.

**Figure 2.**
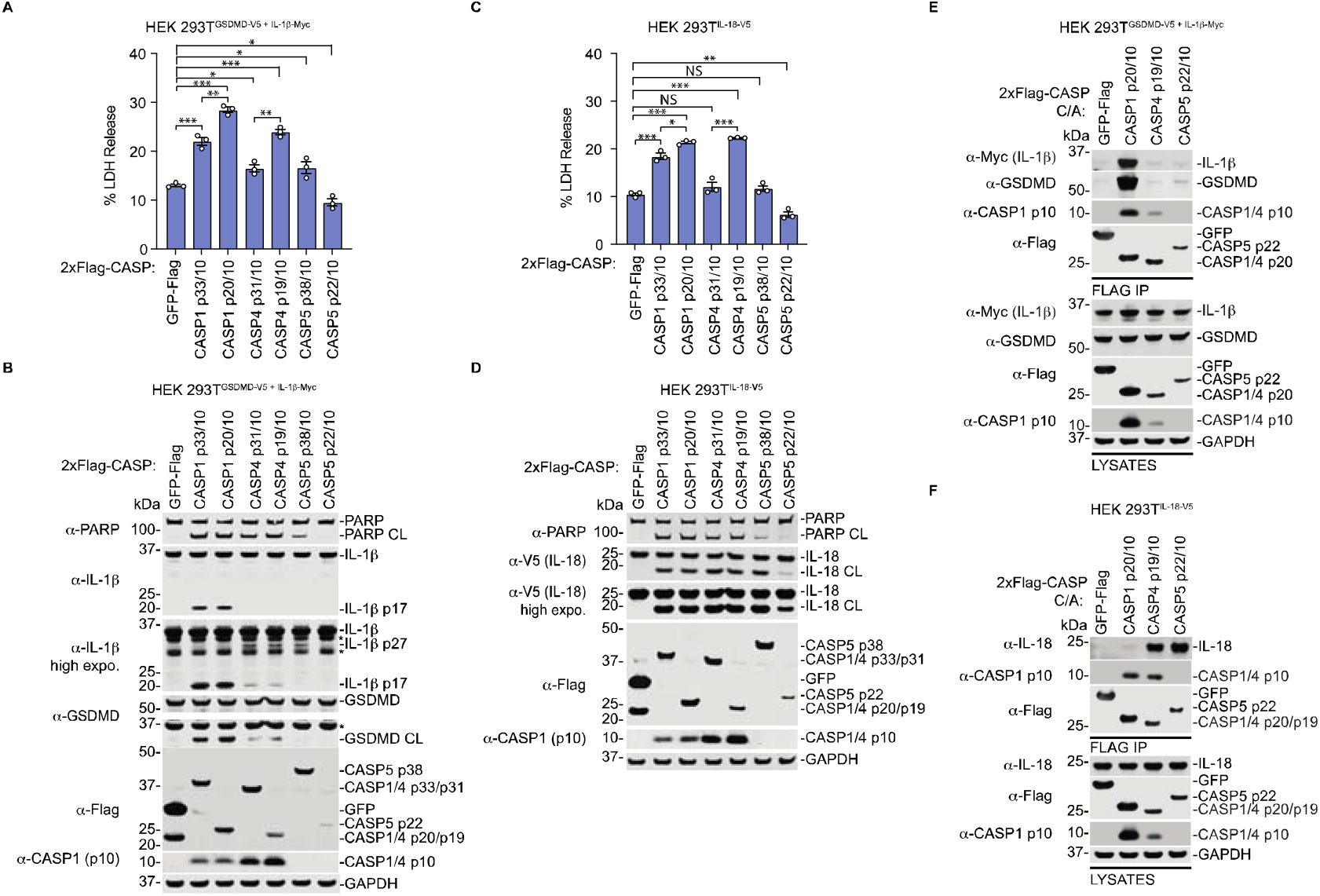
CASP4/5 cleave IL-1β and IL-18. (**A**,**B**) HEK 293T cells stably expressing GSDMD-V5 and IL-1β-Myc were transiently transfected with the indicated constructs. After 24 h, samples were analyzed for LDH release (**A**) and immunoblotting (**B**). (**C**,**D**) HEK 293T cells stably expressing IL-18-V5 were transiently transfected with the indicated constructs. After 24 h, samples were analyzed for LDH release (**C**) and immunoblotting (**D**). (**E**,**F**) HEK 293T cells stably expressing GSDMD-V5 and IL-1β-Myc (**E**) or IL-18-V5 (**F**) were transfected with the indicated catalytically inactive caspase constructs. After 48 h, the cells were harvested and subjected to anti-FLAG IP followed by immunoblot analysis. Data are means ± SEM of three biological replicates. ***P < 0.001, **P < 0.01 and *P < 0.05 by two-sided Student’s *t*-test compared with control. *Represents non-specific bands in immunoblots. Data are representative of three or more independent experiments.

We next expressed the caspase species in HEK 293T cells stably expressing IL-18 and as expected for cells lacking GSDMD^29^, there was PARP processing, indicating that the caspases induced apoptosis in the absence of GSDMD. We note that some PARP processing also occurred in the GSDMD expressing cells, signifying that both apoptosis and pyroptosis pathways can be engaged if there is enough of the functional enzyme species present. Of note, a recent study identified caspase-7 as a substrate of CASP4, implying there may be crosstalk between the non-canonical inflammasome pathway and the apoptotic pathway^30^. There was LDH release in the GSDMD deficient cells due to secondary necrosis but it was less than the LDH released in GSDMD expressing cells (**Fig. 2C**)^5,29,31^. Remarkably, all active caspase species seemed to process IL-18 equivalently (**Fig. 2D**). To gain more insight into the molecular basis of why CASP4/5 did not cleave IL-1β or GSDMD as efficiently as CASP1 but cleaved IL-18 as efficiently, we compared the binding of these substrates to the p20/10 species, which bind inflammatory substrates strongly. In agreement with our previous data, neither CASP4 p19/10 nor CASP5 p22/10 bound GSDMD or IL-1β as strongly as CASP1 p20/10 (**Fig. 2E**). Conversely, more IL-18 was bound to both CASP4 p19/10 and CASP5 p22/10 compared to CASP1 p20/10 (**Fig. 2F**), suggesting that this increased binding compared to CASP1 permits processing to the same degree as CASP1. This data implies that CASP4/5 may preferentially cleave IL-18 before other substrates in cells.

### The P4–P1 tetrapeptide sequence of IL-1β regulates processing by inflammatory caspases

Recent *in vitro* work using recombinant CASP4/5 demonstrated that the identity of the residues adjacent to the caspase cleavage site can influence the processing of substrates^32,33^. While some enhancement in the catalytic efficiency of non-canonical inflammasomes for processing IL-18 was achieved by substituting the P1’–P4’ region with the sequence of GSDMD, these substitutions in IL-1β had limited effect on catalysis^32^. This raises 3 possibilities that are not mutually exclusive: 1) the P4–P1 (rather than the P1’–P4’) region is the major determinant of IL-1β catalysis, 2) the GSDMD sequence is not the optimal sequence for enhancing catalysis, or 3) some other structural features modulate IL-1β recognition and catalysis, such as an exosite. Interestingly, the processing of GSDMD by inflammatory caspases was shown to be sequence independent and instead relied on recognition of an exosite^16^. We wondered if processing of IL-1β was similarly sequence independent. We reasoned that processing of IL-1β at the canonical site (D116) would be sequence dependent since CASP4/5 engaged and processed IL-1β at an alternative site to generate a 27 kDa (IL-1β p27) fragment, but not the p17 fragment (**Fig. 2B**). To test this, we mutated the P4–P1 tetrapeptide sequence of IL-1β to match the tetrapeptide sequence of known caspase substrates (**Fig. 3A**). For example, CASP1 cleaves wildtype IL-1β (IL-**1**β^WT^) at D116, which harbors a YVHD_116_ sequence. We mutated the YVHD sequence to LESD to match the tetrapeptide sequence of IL-18, to generate IL-1β^LESD^. We mutated YVHD to IAND (IL-1β^IAND^) to match the tetrapeptide sequence of IL-1α, which was reported to be processed by CASP5^34^, and generated an IL-1β^AAAD^ mutant which abolished all specificity. All the IL-1β tetrapeptide mutants were HA-tagged at the C-terminus. We transiently co-expressed the IL-1β tetrapeptide mutants with the catalytically active caspase p20/10 constructs in HEK 293T cells expressing GSDMD-V5 and IL-1β^WT^-Myc (**Fig. 3B**). As previously observed, only CASP1 p20/10 significantly processed IL-1β into the p17 fragment (**Fig. 3B**). In support of our hypothesis, we observed that IL-1β^LESD^ was processed by CASP1/4/5 into the active p17 fragment (**Fig. 3B**). Notably, processing of IL-1β^AAAD^ into the p17 fragment was significantly attenuated for all caspases, including CASP1. CASP1 was able to process IL-1β^IAND^ to generate the p17 fragment, but CASP1 also generated a substantial amount of the alternatively processed p27 fragment, suggesting that processing at the canonical site (D116) was suboptimal (**Fig 3B**). Intriguingly, despite CASP5 being reported to cleave IL-1α, CASP5 p22/10 failed to cleave IL-1β^IAND^ to generate the p17 product but rather, generated the p27 product (**Fig. 3B**). These findings suggested that CASP5 may not cleave IL-1α, and indeed, we did not observe processing of wildtype IL-1α by any of the caspases when activated by overexpression or using an inducible system (described below) (**Supplemental Fig. S1 A,B**).

**Figure 3.**
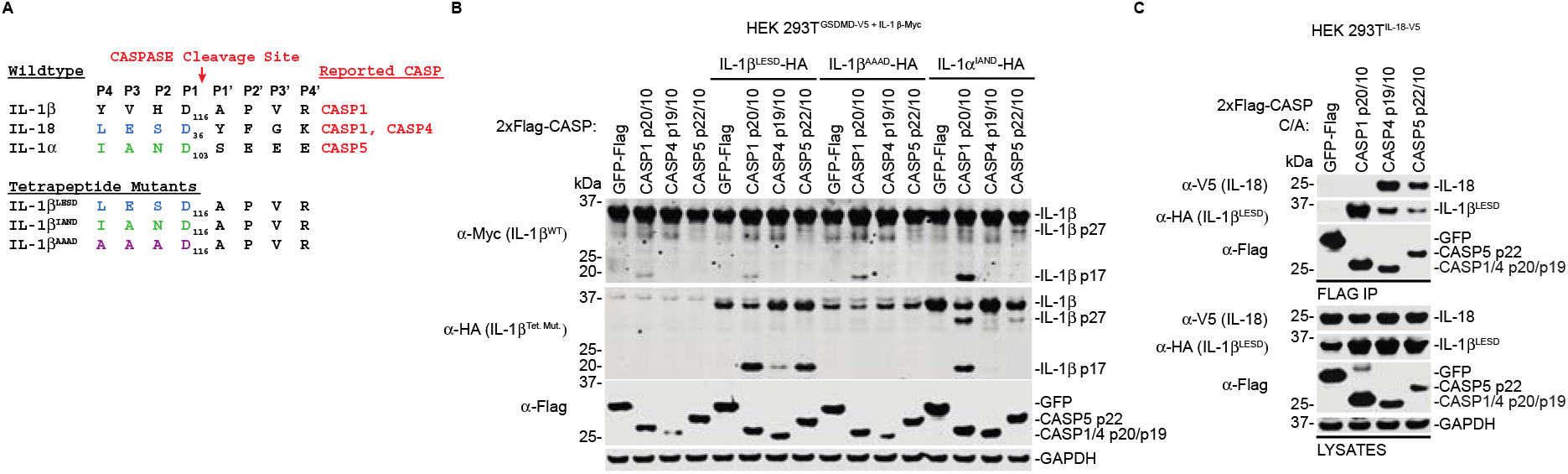
The P4 – P1 tetrapeptide sequence of IL-1β regulates processing by CASP1/4/5. (A)Schematic depicting the sequence of wildtype IL-1β, IL-18, IL-1α, the caspases that were reported to cleave these substrates, and the tetrapeptide mutants. (**B**) HEK 293T cells stably expressing GSDMD-V5 and IL-1β-Myc were transiently co-transfected with the indicated catalytically active caspase constructs and IL-1β tetrapeptide mutants (Tet. Mut.) for 24 h prior to immunoblot analysis. (**C**) HEK 293T cells stably expressing IL-18-V5 were transiently co-transfected with the indicated catalytically inactive (C/A) caspase constructs and the IL-1β mutant, in which the tetrapeptide sequence was substituted for the sequence found in IL-18 (IL-1β^LESD^). 48 h post transfection, samples were subjected to anti-FLAG immunoprecipitation followed by immunoblot analysis. Data are representative of two or more independent experiments.

We next wanted to determine the molecular basis governing the processing of IL-1β^LESD^ by the inflammatory caspases to generate the bioactive p17 fragment but not the other tetrapeptide mutants. Given that wildtype IL-18 binds CASP4/5 strongly, we hypothesized that IL-1β^LESD^ would have increased binding to CASP4/5 compared to IL-1β^WT^. We co-transfected IL-1β^LESD^ with the catalytically inactive caspases and performed anti-FLAG immunoprecipitations to assess binding. Consistent with our hypothesis, we did not detect binding of wildtype IL-1β to CASP4/5 (**Fig. 1D,F**) but we detected binding of IL-1β^LESD^ to CASP4 p19/10 and CASP5 p22/10 (**Fig. 3C**). Taken together, this data suggests that the tetrapeptide motif of IL-1β plays a critical role in regulating its molecular interactions with inflammatory caspases to fine tune inflammation.

### Caspase activation by dimerization is sufficient to cleave inflammatory substrates

The consensus mechanism for initiator caspase activation involves initial dimerization, which allows the caspase to gain basal activity for itself, leading to IDL autoproteolysis and subsequent maturation^35^. Recently, the DmrB dimerization system has been used to induce the dimerization and activation of caspases ^12,13^. Briefly, replacing the CARD domain of the caspase with the DmrB domain enables precise and controlled dimerization and activation in the presence of the small molecule AP20187. To further probe the processing of inflammatory substrates, we generated HEK 293T cells stably expressing either GSDMD-V5 or IL-18-V5 along with either ΔCARD DmrB-CASP1, −4, or −5 (hereafter referred to as DmrB-CASP1/4/5). We treated these cells with AP20187 for 1 hour or 24 hours, and in agreement with our previous data, we observed GSDMD (**Fig. 4A**) and IL-18 (**Fig. 4B**) processing. Although there was some GSDMD processing within 1 h, we did not detect LDH release within this timeframe, but LDH release was observed at 24 hours. Of note, DmrB-CASP1 exhibited the highest LDH release in both cell lines (**Fig. 4A,B**) and was the only one that induced apoptosis that progressed to secondary necrosis in cells expressing IL-18 (**Fig. 4B**). In agreement with the LDH release and our prior data, DmrB-CASP1 had more GSDMD processing than DmrB-CASP4 and DmrB-CASP5. However, DmrB-CASP5 processed IL-18 nearly to the same degree as DmrB-CASP1 within 1 hour, and all caspases processed IL-18 after 24 hours (**Fig. 4B**). We next assessed the processing of IL-1β and the tetrapeptide mutants using the DmrB-caspases. We transiently transfected wildtype or the IL-1β tetrapeptide mutants into HEK 293T cells expressing GSDMD-V5 and DmrB-CASP1, −4, or −5, then treated with AP20187 for 24 hours (**Fig. 4C**). Only Dmrb-CASP1 cleaved IL-1β^WT^ at the canonical site to yield the p17 fragment. However, IL-1β^LESD^ was equivalently processed by DmrB-CASP1/4/5, and none of the caspases cleaved IL-1β^AAAD^ or IL-1β^IAND^ (**Fig. 4C**). This orthogonal method of caspase activation confirms that CASP4/5 do not process GSDMD as efficiently as CASP1, but IL-18 maturation occurs nearly as efficiently with CASP4/5 as with CASP1. Furthermore, these results reaffirm that the P4-P1 tetrapeptide motif of IL-1β regulates processing by inflammatory caspases.

**Figure 4.**
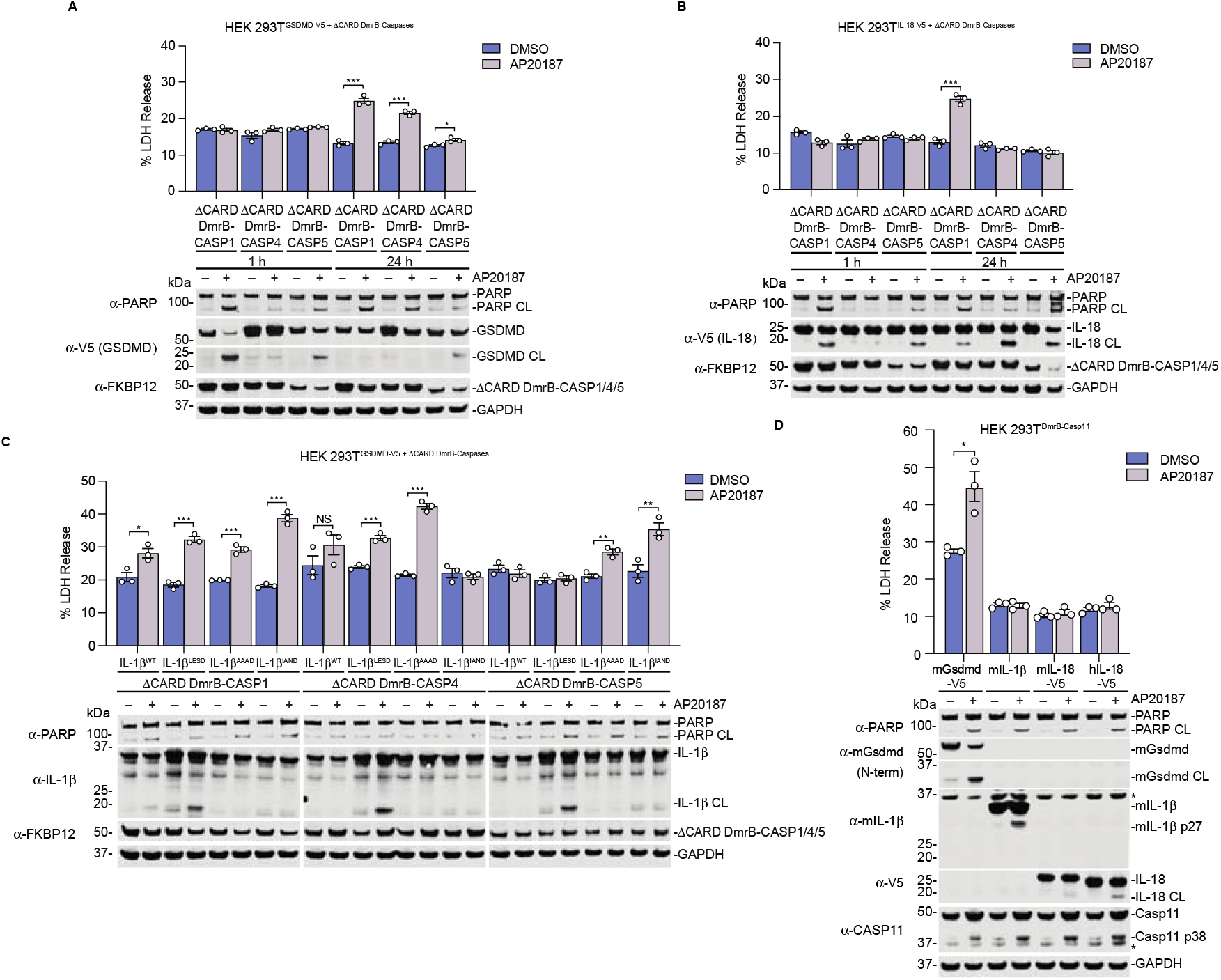
Dimerization of ΔCARD DmrB-CASP1/4/5/11 mediates processing of inflammatory substrates. (**A**,**B**) HEK 293T Cells stably expressing ΔCARD DmrB-CASP1, −4, or −5 and GSDMD-V5 (**A**) or IL-18-V5 (**B**) were treated with AP20187 (1 μM) for 1 h or 24 h. Cell death was measured by LDH and samples were analyzed by immunoblotting. (**C**) HEK 293T cells stably expressing ΔCARD DmrB-CASP1, −4, or −5 and GSDMD-V5 were transiently transfected with the indicated IL-1β constructs for 24 h before the addition of AP20187 (1 μM) for 24 h. Cell death was measured by LDH and samples were analyzed by immunoblotting. (**D**) HEK 293T cells stably expressing ΔCARD Dmrb-CASP11 were transiently transfected with the indicated constructs. After 24 h, samples were treated with AP20187 (1 μM) for 24 h then analyzed for LDH release and immunoblotting. Data are means ± SEM of three biological replicates. ***P < 0.001, **P < 0.01 and *P < 0.05 by two-sided Student’s *t*-test compared with control. *Represents non-specific bands in immunoblot. Data are representative of two or more independent experiments.

CASP11 is considered the mouse ortholog of human CASP4/5. We wondered if there were differences between the mouse and human non-canonical inflammasomes in their ability to process cytokines. To address this, we generated HEK 293T cells stably expressing ΔCARD DmrB-CASP11 and transiently transfected mouse Gsdmd (mGsdmd), IL-1β (mIL-1β), IL-18 (mIL-18), or human IL-18 (hIL-18) into the cells. After 24 h post transfection, DmrB-CASP11 was activated with AP20187 for 24 h then samples were analyzed for LDH release and immunoblotting (**Fig. 4D**). As expected, the addition of AP20187 induced pyroptosis in the mGsdmd transfected cells, as evidenced by LDH release and mGsdmd processing into the pyroptosis-inducing p30 fragment (**Fig. 4D**). Notably, DmrB-CASP11 induced mIL-1β processing at an alternative site to yield the deactivated p27 fragment, but not the active p17 fragment (**Fig. 4D**). Unlike CASP4/5, DmrB-CASP11 failed to significantly process IL-18 into the bioactive mature species (**Fig. 4D**), suggesting a functional divergence between mouse and human non-canonical inflammasomes.

### LPS activated non-canonical inflammasomes cleave IL-1β and IL-18

The non-canonical inflammasomes are activated by intracellular LPS^17–20^. We next sought to determine if activation by the native ligand (LPS) would induce cytokine processing. We transiently transfected LPS into HEK 293T cells expressing CASP4/5 or 2xFLAG-CASP4/5 and IL-18. We detected processing of IL-18 into mature IL-18 in LPS-transfected cells (**Fig. 5A**). Moreover, the presence of the N-terminal 2x-FLAG tag did not affect the activity of CASP4/5 as the maturation of IL-18 was comparable to that of the untagged CASP4/5-expressing cells (**Fig. 5A**).

**Figure 5.**
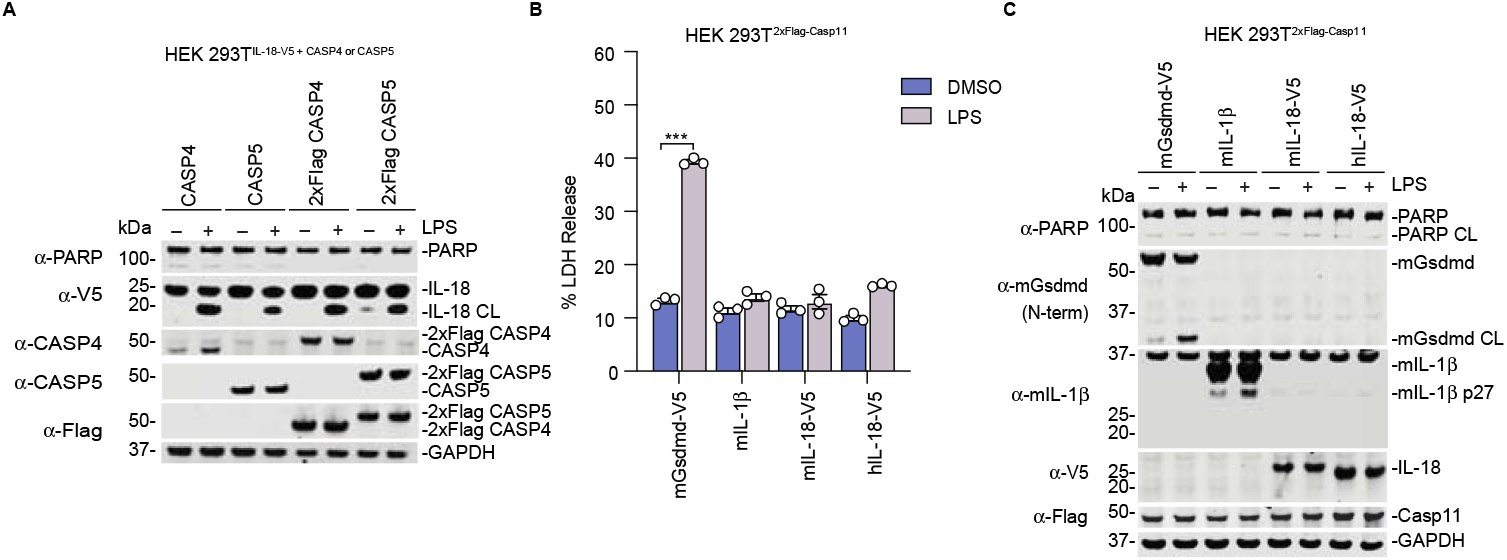
LPS activated non-canonical inflammasomes cleave IL-1β and IL-18. (**A**) HEK 293T cells stably expressing IL-18-V5 and either CASP4 or CASP5 were transfected with LPS (25 μg/mL) for 24 h before samples were collected and analyzed by immunoblotting. (**B**,**C**) HEK 293T cells stably expressing 2xFLAG-CASP11 were transiently transfected with the indicated constructs. After 24 h, samples were transfected with LPS (25 μg/mL) for 24 h then analyzed for LDH release (**B**) and immunoblotting (**C**). Data are means ± SEM of three biological replicates. ***P < 0.001, **P < 0.01 and *P < 0.05 by two-sided Student’s *t*-test compared with control. Data are representative of two or more independent experiments. *Represents non-specific bands in immunoblot. The small m or h represents mouse or human versions of the proteins respectively.

Our results with DmrB-CASP11 indicate that CASP11 cleaves IL-1β but not IL-18. We wanted to determine if LPS-activated CASP11 would similarly cleave IL-1β to generate the p27 fragment, but not cleave IL-18. We transiently transfected the inflammatory substrates into HEK 293T cells stably expressing 2x-FLAG CASP11 and activated with LPS. Analogous to the DmrB-CASP11 results, LPS activated CASP11 induced pyroptosis, as determined by GSDMD processing and LDH release (**Fig. 5B,C**). Importantly, LPS-activated CASP11 processed mIL-1β to yield the p27 fragment, but not the active p17 fragment, and failed to process IL-18 (**Fig. 5C**). Altogether, this data implies that the inactivation of IL-1β signaling is conserved between humans and mice, but the activation of IL-18 is not.

### Non-canonical inflammasome activation induces IL-1β and IL-18 processing in human macrophages and epithelial cells

The non-canonical inflammasomes are expressed in both myeloid and epithelial cells and have a well-documented function in playing an important role in mediating host protection against invading pathogens^17,19,24–26^. While epithelial cells express IL-18, there is limited IL-1β expression even with toll-like receptor (TLR) stimulation^26,36,37^. On the other hand, monocytes and macrophages robustly express IL-1β upon TLR stimulation, which can occur during infection with gram negative bacteria^36^. To gain insight into the function of non-canonical inflammasomes in different cell types, we sought to determine the activity of CASP4/5 in more physiologically relevant cells. We first differentiated THP1 cells into macrophages, primed with LPS to ensure IL-1β expression, then transfected with LPS to induce endogenous CASP4/5 activation. As anticipated, LPS induced significant LDH release in control and *CASP1* knockout (KO) cells, indicating CASP4/5-mediated pyroptosis (**Fig. 6A**). Indeed, LPS-transfected THP1 macrophages displayed GSDMD processing in both control and *CASP1* KO cells (**Fig. 6B**). To delineate the contribution of CASP4/5 to IL-1β and IL-18 processing in macrophages, we treated the cells with a broad-spectrum pharmacological inhibitor (ZVAD) that targets all caspases, and an inhibitor that is more specific for CASP1 (VX765)^38,39^. We then transfected LPS to activate CASP4/5 and 24 hours later, separated the supernatants and lysates for analysis by immunoblotting. During pyroptosis, inflammatory substrates are released into the supernatant through the gasdermin pores, so we expected to see the processed substrates in the supernatants. In accord with this, activation of CASP4/5 in control cells led to robust processing of IL-1β into the p27 fragment, which was nearly completely abrogated by ZVAD, and significantly attenuated by VX765 (**Fig. 6B**). NLRP3 is activated downstream of non-canonical inflammasomes, which activates CASP1 to mediate IL-1β processing into the p17 fragment^22,23^. Consistent with this, we also observed IL-1β maturation into the bioactive p17 fragment in control cells, which was completely abrogated by both inhibitors (**Fig. 6B**).

**Figure 6.**
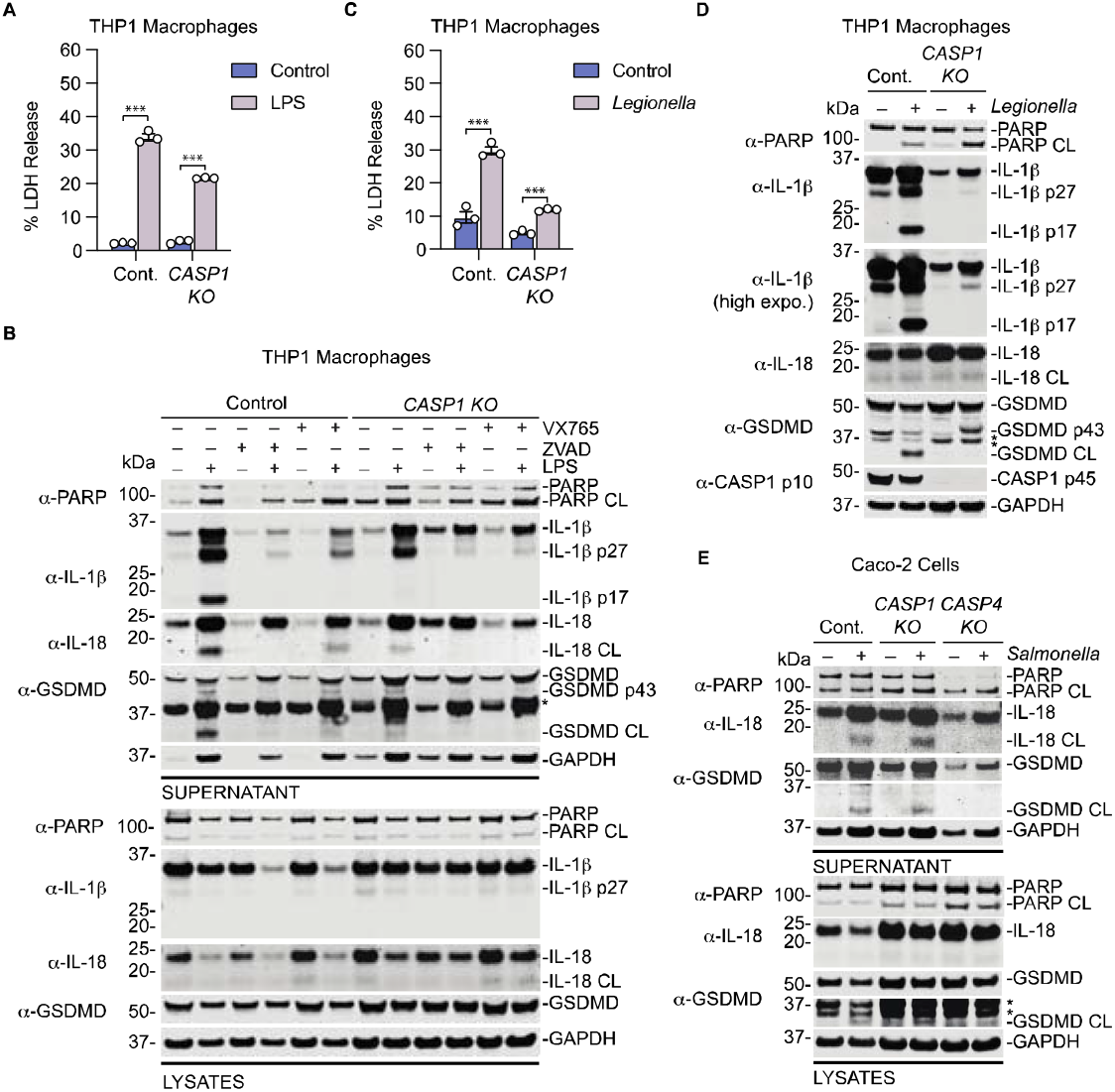
Cytosolic LPS and pathogenic infections induce non-canonical inflammasome mediated processing of inflammatory substrates in human macrophages and epithelial cells. (**A**,**B**) THP1 cells were terminally differentiated into macrophages with phorbol 12-myristate 12-acetate (50 ng/mL) for 24 h and primed with LPS (5 μg/mL) for another 24 h. Where indicated, THP1 macrophages were treated with ZVAD (40 μM) or VX765 (40 μM) 30 minutes before LPS transfections. Cells were then transfected with LPS (25 μg/mL). 24 h after LPS transfection, samples were analyzed for LDH release (**A**) and immunoblotting (**B**). (**C**,**D**) THP1 cells were terminally differentiated into macrophages with phorbol 12-myristate 12-acetate (40 ng/mL) for 48 h. Cells were then treated with *Legionella pneumophila* (MOI = 20) for 7 h before the supernatants were analyzed for LDH release (**C**) and then combined with the lysates for immunoblotting (**D**). € Caco-2 cells were primed with 100 ng/ml Pam3CSK4 for 3 h, then infected with *Salmonella* Typhimurium (MOI = 60) for 6 h. Cells and their supernatants were collected separately, samples were precipitated, and analyzed by immunoblotting. Data are means ± SEM of three biological replicates. ***P < 0.001 and **P < 0.01 by two-sided Student’s *t*-test compared with control. *Represents non-specific bands in immunoblot. Data are representative of two or more independent experiments.

Importantly, we observed processing of IL-1β into the p27 fragment in *CASP1* KO cells, indicating that this processing event results from CASP4/5 activity (**Fig. 6B**). We did not detect the presence of the mature IL-1β p17 fragment in *CASP1* KO cells, suggesting CASP4/5 do not induce significant processing of IL-1β at the canonical site in human macrophages (**Fig. 6B**). Critically, the appearance of the p27 fragment was abrogated in the presence of caspase inhibitors in *CASP1* KO cells (**Fig. 6B**). It should be noted that pharmacological inhibitors of caspases can cross-react with other caspases and the CASP1 inhibitor VX765 also inhibits CASP4/5. Similar findings were noted for the release of mature IL-18 into the supernatant. We observed CASP1-independent release of mature IL-18, which was abrogated by ZVAD and VX765, signifying that CASP4/5 cleave IL-18 to generate the bioactive species in human macrophages (**Fig. 6B**).

We next wanted to assess whether CASP4/5 mediate processing of IL-1β and IL-18 in the context of a natural bacterial infection. We therefore infected THP1 macrophages with *Legionella pneumophila* (hereafter called *Legionella*) and like LPS, *Legionella* infection resulted in increased LDH release in control and *CASP1* KO cells compared to uninfected cells (**Fig. 6C**). *Legionella* infection induced the processing of IL-1β into both the p27 and p17 fragments in control cells, but only the p27 fragment was generated in *CASP1* KO cells (**Fig. 6D**). We detected a slight increase in IL-18 processing in *Legionella* infected cells. Collectively, our data indicates that CASP4/5 directly process IL-1β and IL-18 in macrophages during bacterial infection.

Because prior studies have implicated CASP4/11 in IL-18 processing in epithelial cells, we wanted to determine the role of non-canonical inflammasome activation on IL-18 processing in epithelial cells. We thus infected Caco-2 cells with *Salmonella* Typhimurium, which was previously documented to activate CASP4 in Caco-2 cells^25,26^. Like CASP4/5 activation in macrophages, *Salmonella* infection induced GSDMD processing and IL-18 maturation and release into the supernatants (**Fig. 6E)**. As previously reported, IL-18 processing was CASP1-independent and CASP4-dependent (**Fig. 6E)**^25,26^. Taken together, our data suggests that IL-18 is a direct substrate for the non-canonical inflammasomes. However, in certain cell types that express IL-1β, non-canonical inflammasome activation leads to IL-1β processing at an alternative site to generate a p27 fragment that was previously reported to be functionally inactive^28^.

### CASP4/5 process IL-1β at D27 and IL-18 at D36

CASP1 is known to process IL-1β at D27 to generate a fragment that is ∼27 kDa^40,41^. We sought to identify the alternative processing site that gives rise to IL-1β p27. We postulated that CASP1 prefers to cleave IL-1β at D116 to generate IL-1β p17, but CASP4/5 preferentially cleave IL-1β at D27 to generate IL-1β p27. To test this hypothesis, we mutated D27 to alanine and co-transfected IL-1β D27A (IL-1β^D27A^) with the catalytically active species of CASP1/4/5 into HEK 293T cells. As anticipated, CASP1 p33/10 and p20/10 processed IL-1β^WT^ and IL-1β^D27A^ to generate IL-1β p17 whereas CASP4 p31/10 and p19/10 primarily processed IL-1β^WT^ to generate the p27, and modestly generated the p17 fragment (**Fig. 7A**). Notably, CASP4 failed to cleave IL-1β^D27A^ to generate IL-1β p27, demonstrating that D27 is indeed the processing site for CASP4 (**Fig. 7A**). In agreement with our prior data, CASP5 processed IL-1β^WT^ and generated only the p27 fragment, which was abolished in IL-1β^D27A^ transfected cells (**Fig. 7A**). We also discovered that CASP5 cleaves IL-18 and wanted to confirm the site of processing. Because CASP1/4 process IL-18 at D36^40,41^ and CASP5 processing of IL-18 generated a fragment that was the same size as that of CASP1/4 processed IL-18, we hypothesized that CASP5 processed IL-18 at D36. Indeed, CASP1/4/5 processed IL-18^WT^ to generate the mature species, which was abolished in IL-18^D36A^ transfected cells (**Fig. 7B**). Thus, our data reveals that the non-canonical inflammasomes process IL-1β at D27 and IL-18 at D36, establishing IL-1β and IL-18 as bona fide substrates of the non-canonical inflammasomes (**Fig. 7C**).

**Figure 7.**
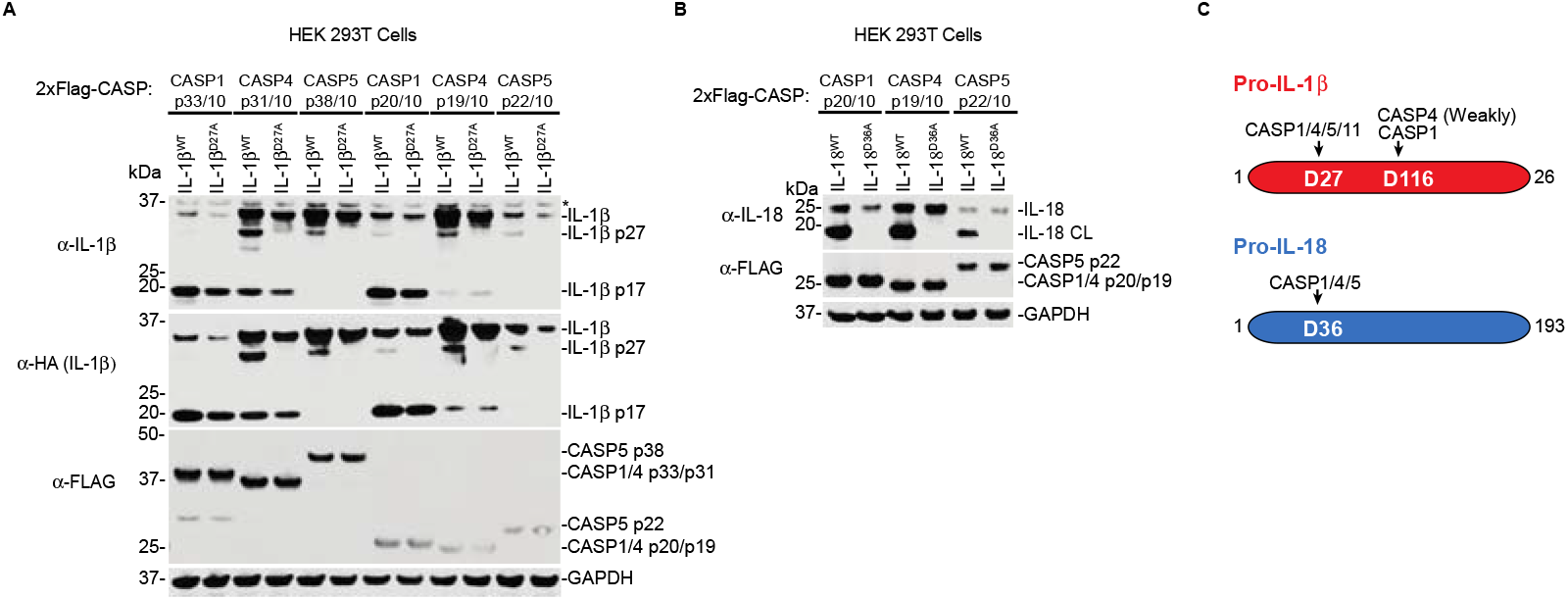
CASP4/5 preferentially cleave IL-1β at D27 and IL-18 at D36. (**A**,**B**) HEK 293T cells were transiently co-transfected with the indicated constructs. After 24 h, samples were harvested and analyzed by immunoblotting. (**C**) Schematic of inflammatory substrates and representative cleavage sites by inflammatory caspases. Data are representative of three or more independent experiments. *Represents non-specific bands in immunoblot.

## Discussion

All inflammatory caspases are known to cleave the effector protein GSDMD to induce pyroptosis, but whether these caspases process GSDMD to the same degree when activated in cells has remained unclear because ligand-mediated activation of inflammatory caspases proceed with different kinetics. For example, lethal factor (LF) and the small molecule drug Val-boroPro (VbP) are both activators of the NLRP1B inflammasome, but LF-induced pyroptosis occurs faster than VbP-induced pyroptosis^42,43^. In this study, we developed an expression system that allowed us to compare the interactions of specific caspase species with their putative substrates as well as evaluate the activity in cells independent of the kinetics of ligand-mediated activation. Furthermore, we utilized an orthogonal chemical biology approach to inducibly dimerize and activate the inflammatory caspases that corroborates the findings from our expression assays. Importantly, these findings are congruent with results obtained from activating the inflammatory caspases using native ligands and pathogens in physiologically relevant cells.

Recent studies have reported that the dominant active caspase-1 species in cells is a transient p33/10 species, and further processing to the p20/10 species generates an unstable protein that rapidly loses protease activity^9^. Consistent with these findings, several studies have demonstrated that interdomain linker processing is critical for pyroptosis^9,12–14^. Interestingly, structural studies demonstrate that the p20/10 species of the inflammatory caspases utilize an exosite to bind to GSDMD^15,16^. However, the structural basis of how inflammatory caspases bind to other substrates such as IL-1β and IL-18 are not fully defined. Thus, we sought to characterize the inflammatory caspase species and their Interactions with the inflammatory substrates, GSDMD, IL-1β, and IL-18. Our data suggests that the p20/10 species of CASP1/4/5 are the species that bind strongly to the inflammatory substrates (**Fig. 1**). However, some notable differences exist. For example, CASP1 p20/10 binds strongly to GSDMD and IL-1β, and weakly to IL-18. In contrast, CASP4/5 p20/10 binds strongly to IL-18, and weakly to GSDMD and IL-1β. Although the p35/10 species displayed weak interactions with the inflammatory substrates, our data suggests that the weak interactions are sufficient to mediate processing of these substrates (**Fig. 2**). We note that for CASP1, less of the p33/10 was pulled down in the IP samples and this may account for why there appears to be less binding of the substrates (**Fig. 1**). However, it is worth mentioning that less CASP4 p19/10 was pulled down compared to CASP1 p20/10, but more IL-18 was bound to CASP4 p19/10, suggesting that if a functional enzyme complex is formed, strongly bound substrates would co-IP with the caspase species (**Fig. 2F**). Indeed, a functional CASP1 p33/10 enzyme complex was formed as evidenced by the processing of substrates such as IL-18 and IL-1β and the presence of the p10 in the IP samples (**Fig. 1**,**2**). If CASP1 p33/10 can mediate cytokine processing, then a major question is why is ASC required? ASC acts as a signal amplifier that facilitates full autoproteolysis of pro-CASP1 into the p20/10 species,^10,11,44^ precisely the species that binds strongly to the cytokines. ASC may have additional functions that are important for IL-1β processing, but we postulate that in part, it helps generate enough of the short-lived p20/10 species that recruits and processes IL-1β. These differences in substrate binding capabilities likely help to regulate inflammation by the inflammatory caspases under different pathophysiological conditions. Future studies will help uncover the role of ASC in cytokine maturation.

The non-canonical inflammasomes are critical for host responses to invading gram negative bacteria. When activated, CASP4/5 in humans and CASP11 in mice, are known to cleave GSDMD to mediate pyroptosis, but whether other direct substrates of the non-canonical inflammasomes exist has remained unclear. Here, we discovered that CASP4/5 directly process IL-18 at D36 to generate the bioactive species. Notably, IL-18 is a potent activator of natural killer and T cells,^45^ thus, processing by CASP4/5 during infection likely confers an advantage for mounting an immune response. These findings are in line with recent genetic evidence that CASP4 is required for IL-18 maturation and secretion in epithelial cells during bacterial infection^24–26^. Furthermore, our data indicates that the ability of CASP4/5 to process IL-18 is not limited to just epithelial cells, but spans across other cell types, including macrophages (**Fig. 6**). Intriguingly, while human CASP4/5 robustly process IL-18, the mouse orthologue of CASP4/5, CASP11, is unable to process IL-18 in cells, consistent with *in vitro* reports^32,33^. Recent studies also suggest CASP11 does not process IL-18 in response to non-canonical agonists,^46^ implying a functional divergence between humans and mice with regard to IL-18 processing. The impact of this divergence on host responses to bacteria warrants future investigation.

Unexpectedly, although we did not detect binding of IL-1β to CASP4/5, we could detect processing by both CASP4/5 to generate a p27 fragment (Fig. 2,6,7). Remarkably, the p27 fragment of IL-1β was the dominant species generated when CASP4/5 were activated in THP1 macrophages by LPS transfection or with bacterial infection (**Fig. 6**). During the preparation of this manuscript, another group used DmrB-CASP4 and reported processing of IL-1β to generate the mature p17 fragment^47^. We also detected modest processing of IL-1β by CASP4 to generate the p17 fragment, which was significantly less than the p17 fragment generated by CASP1 in HEK 293T cells (**Fig. 2**). However, we could not detect IL-1β p17 when CASP4/5 were activated in human macrophages, suggesting this may not be physiologically relevant (**Fig. 6**). It is possible that the processed p17 fragment generated by CASP4/5 activity was below the limit of detection in our macrophage assays. Regardless, the fact that we could detect CASP4/5-mediated processing of IL-1β to the p27 fragment suggests it is the more physiologically relevant processing event. CASP11 also processed mouse IL-1β to generate a p27 fragment (**Fig. 5**), implying that the ability to generate IL-1β p27 is at least evolutionarily conserved between mice and humans – unlike IL-18 processing.

The functional impact of non-canonical inflammasome processing of IL-1β into the p27 fragment remains to be determined. It is tempting to speculate that at some point during evolution, IL-1β signaling was detrimental to the host, and the non-canonical inflammasomes evolved to deactivate IL-1β signaling. In support of this hypothesis, IL-1β blocking antibodies were recently demonstrated to significantly reduce the incidence of lung cancer^48^. Notably, pro-inflammatory cytokines, including IL-1β and IL-18, contribute to the excessive inflammation and pathogenesis of sepsis that leads to organ failure, and blocking these cytokines confers protection in acute animal models^49^. Also in support of the notion that the non-canonical inflammasomes may have evolved to attenuate excessive inflammation downstream of bacterial infections, recent ancestral reconstruction revealed that CASP4/5 evolved after CASP1 from an ancestral caspase that cleaves IL-1β to yield the p27 fragment, which was reported to not signal to the IL-1 receptor^28^. We mapped out the processing site that generates IL-1β p27 by the non-canonical inflammasomes to D27 (**Fig. 7**). However, whether CASP1 can further process IL-1β p27 to generate the mature, active p17 fragment remains unknown. Fascinatingly, extant Carnivora harbor a CASP4 enzyme scaffold that has CASP1-like substrate specificity^50^. Perhaps in the Carnivora, IL-1β p17 was detrimental but later became essential for survival, and thus, Carnivora CASP4 gained CASP1-like activity. Future studies are needed to delineate the biological contributions of IL-1β p27 and p17 to host defense or regulating inflammation during bacterial infection.

GSDMD is recognized by inflammatory caspases via an exosite, and the processing is sequence independent^16^. In stark contrast, we discovered that the processing of IL-1β is sequence dependent. When we substituted the native tetrapeptide sequence of IL-1β for that of tetrapeptide sequences from other inflammatory caspase substrates, this conferred processing specificity to that corresponding caspase. Specifically, we substituted the tetrapeptide sequence of IL-1β (YVHD) for that of IL-18 (LESD), which we demonstrated was processed by CASP4/5, the mutant IL-1β was then processed by CASP4/5 at that tetrapeptide site (**Fig. 3)**. However, if YVHD is replaced by AAAD, a sequence that has no specificity for a particular caspase, in contrast to GSDMD, this attenuates processing at that site. Rather than processing at that site to generate the bioactive p17 fragment, IL-1β^AAAD^ was processed at D27 to yield a 27 kDa fragment that is predicted to be inactive (**Fig. 3**). This reengineering of IL-1β substrate specificity led us to discover that CASP5, which is reported to cleave IL-1α ^34^, does not process IL-1α (**Fig. S1**). In fact, we did not observe IL-1α processing by CASP1/4/5 when activated via different mechanisms. This is consistent with past and present studies that failed to detect a role for inflammatory caspases in processing IL-1α, but instead identified granzyme B and calpains as the proteases that mediate IL-1α processing^51,52^. The incongruent findings could arise from the use of different cell types and *in vitro* vs in cell assays. Altogether, these findings indicate that the tetrapeptide sequence in IL-1β regulates IL-1β recruitment and processing by CASP1/4/5. We postulate that peptide inhibitors based on the LESD scaffold from IL-18 may potentially serve as selective inhibitors of CASP4/5 over CASP1. Future structural studies are necessary to dissect the molecular basis of distinct IL-1β and IL-18 recognition and processing by inflammatory caspases.

In summary, report two new direct substrates (IL-1β and IL-18) for the non-canonical inflammasomes. CASP4/5 cleave IL-18 at D36 to generate the bioactive fragment, but process IL-1β at D27 to yield an inactive p27 fragment (**Fig. 7C**). Additionally, we discovered that CASP11 similarly processes mIL-1β to generate IL-1β p27 but does not process mIL-18. Finally, we found that none of the inflammatory caspases process IL-1α. Hence, the present study offers mechanistic insight into substrate specificities that could help design new therapies for inflammatory disorders and aid our understanding of host responses to inflammation and bacterial infections.

## Materials and Methods

### Antibodies and reagents

Antibodies used include: GSDMD Rabbit polyclonal Ab (Novus Biologicals, NBP2-33422), FLAG® M2 monoclonal Ab (Sigma, F3165), GAPDH Rabbit monoclonal Ab (Cell Signaling Tech, 14C10), CASP1 p20 Rabbit polyclonal Ab (Cell Signaling Tech, 2225s), CASP1 p12/10 Rabbit monoclonal Ab (Abcam, ab179515), CASP4 Rabbit polyclonal Ab (Cell Signaling Tech, 4450S), CASP5 Rabbit monoclonal Ab (Cell Signaling Tech, 46680S), Myc Mouse monoclonal Ab (Cell Signaling Tech, 2276S), V5 Rabbit monoclonal Ab (Cell Signaling Tech, 13202S), hIL-1β Goat Polyclonal Ab (R&D systems, AF-201-NA), hIL-18 Goat Ab (R&D systems, af2548), hIL-1α Recombinant Ab (PeproTech, 200-01A), PARP Rabbit polyclonal Ab (Cell Signaling Tech, 9542S), HA Rabbit monoclonal Ab (Cell Signaling Tech, 3724S), mIL-1β Goat polyclonal Ab (R&D systems, AF-401-NA), CASP11 Rat monoclonal Ab (Novus Biologicals, NB120-10454), FKBP12 Rabbit polyclonal Ab (Abcam, ab24373). IRDye 800CW anti-rabbit (LICOR, 925-32211), IRDye 800CW anti-mouse (LI-COR, 925-32210), IRDye 680CW anti-rabbit (LI-COR, 925-68073), IRDye 680CW anti-mouse (LI-COR, 925-68072). Other reagents used include: LPS-EB Ultrapure (Invivogen, tlrl-3pelps), VX-765 (Apexbio Technology LLC, 50-101-3604), Z-VAD-FMK (Enzo Life Sciences, NC9471015), FuGENE HD (Promega, E2311), AP20187 (Tocris™ 6297/5), NP-40 Lysis Buffer Low Salt (Thomas Scientific, C994H79).

### Cell Culture

HEK 293T cells, HeLa cells, Caco-2 cells, and THP1 cells were purchased from ATCC. Caco-2 and THP1 knockout cell lines were previously reported^25,29^. HEK 293T and HeLa cells were cultured in Dulbecco’s Modified Eagle’s Medium (DMEM) with L-glutamine and 10% fetal bovine serum (FBS). THP-1 cells were cultured in Roswell Park Memorial Institute (RPMI) medium 1640 with L-glutamine and 10% fetal bovine serum (FBS). Caco-2 cells (HTB-27; American Type Culture Collection) were maintained in DMEM supplemented with 10% FBS, 100 IU/mL penicillin and 100 mg/mL streptomycin. All cells were grown at 37 °C in a 5% CO_2_ atmosphere incubator.

### Generation of stable cell lines

For generating HEK 293T cells ectopically expressing GSDMD, IL-1β, IL-1α, IL-18, and caspase constructs, plasmids encoding those proteins were packaged into lentivirus by transfecting the vectors (2 μg) along with psPAX2 (2 μg), and pMD2.G (1 μg) using Fugene HD transfection reagent (Promega) into HEK 293T cells. After 2 days, the supernatants were filtered using a 0.45 μm filter, then used to infect HEK 293T cells. After 48 hours, the cells expressing the indicated constructs were selected with hygromycin (200 μg/mL), blasticidin (10 μg/mL), or puromycin (1 μg/mL).

### Cloning

All plasmids were cloned using Gateway technology as previously described^12,29,53^. DNA encoding the indicated proteins were inserted between the *attR* recombination sites and shuttled into modified pLEX_307 vectors (Addgene) using Gateway technology (Thermo Fisher Scientific) according to the manufacturer’s instructions. Proteins expressed from these modified vectors contain an N-terminal *att*B1 linker (GSTSLYKKAGFAT) after any N-terminal tag (2xFLAG) or protein (DmrB) or a C-terminal *att*B2 linker (DPAFLYKVVDI) preceding any C-terminal tag such as V5 or HA. An internal ribosome entry site (IRES) was cloned between the large and small subunits of the caspases to allow separate expression of the two polypeptides. ΔCARD DmrB-CASP1 (residues 92 – 402), ΔCARD DmrB-CASP4 (residues 67– 377), ΔCARD DmrB-CASP5 (residues 128 – 434), and ΔCARD DmrB-CASP11 (residues 71 – 373) were all cloned into a modified pLEX_307 vector and contained the N-terminal *att*B1 linker between the DmrB and caspase sequences. Point mutations were generated using the QuikChange II site-directed mutagenesis kit (Agilent) according to the manufacturer’s instructions.

### Transient transfections

HEK 293T cells were seeded in 12-well culture plates at 2.5 × 10^5^ cells/well in DMEM. The following day, the indicated plasmids were mixed with a control RFP vector to a total of 1.0 μg DNA in 65 μL Opti-MEM and transfected using FuGENE HD (Promega) according to the manufacturer’s protocol. Some samples were further treated (as indicated in each figure legend) with AP20187 (1 μM) or LPS (25 μg/mL)/FuGENE (0.5%). The cells were harvested at the indicated times and analyzed by LDH cytotoxicity and immunoblotting assays as described below.

### LDH cytotoxicity and immunoblotting assays

Supernatants were harvested for LDH analyses at time points indicated and analyzed using the Cyquant LDH Cytotoxicity Assay (Thermo Scientific) according to the manufacturer’s protocol. LDH activity was quantified relative to a lysis control where cells were lysed using NP-40 for 30 minutes. For immunoblotting, protein concentrations were normalized using the DC Protein Assay Kit (Bio-Rad), separated by SDS-PAGE, transferred onto Nitrocellulose membranes (Bio-Rad), and visualized using the Odyssey M Imaging System (LI-COR Biosciences).

### FLAG immunoprecipitations

HEK 293T cells were seeded in 6-well plates at 5 × 10^5^ cells/well in DMEM for 24 h. The cells were then transiently transfected with the indicated constructs. After 48 h, the cells were harvested and lysed by sonication. Lysates were clarified by centrifugation at 21,000 x g for 5 minutes. The soluble fractions were then normalized using the DC Protein Assay (BioRad). 100 μL of the lysates were combined with 100 μL of 2x sample loading buffer and incubated at 95 °C for 10 minutes. Equal protein amounts of the remaining sample lysates were loaded onto Pierce Micro-Spin Columns (Thermo Scientific) containing 100 μL of anti-FLAG-M2 agarose resin (Sigma) and the samples were rotated end-over-end at 4°C overnight. The samples were then washed 3x with 1 column volume (350 μL) of PBS. Proteins were eluted by rotating the resin at room temperature for 1 hour in 100 μL of PBS containing 150 ng/ μL 3×-FLAG peptide (Sigma Aldrich). 100 μL of 2x sample loading buffer was added to the eluate and samples were boiled at 95 °C for 10 minutes. Both lysates and eluates were analyzed by immunoblotting

### LPS transfections of THP-1 cells

THP-1 cells were resuspended in RPMI medium containing 50 ng/ml phorbol 12-myristate 12-acetate (PMA). Cells were plated in 12-well plates at a density of 8 × 10^5^ cells/well. After 24 hours, the media was replaced with fresh RPMI/10% FBS containing 5 μg/ml LPS and the cells were allowed to grow for another 24 h. The media was then replaced with Opti-MEM (0.5 mls/well). Where indicated, cells were treated with ZVAD (40 μM) or VX765 (40 μM) 30 minutes before *LPS transfection. The LPS solution was prepared by adding LPS (25 μg/mL final concentration) and FuGENE (0*.*5% final concentration) to Opti-MEM. This solution was gently mixed by flicking and incubated for 30 mins at room temperature before drop-wise addition to each well*. The supernatants were collected, and cells were lysed by sonication 24 h post transfection. Both supernatants and lysates were precipitated by chloroform/methanol and analyzed by immunoblotting.

### Bacterial Infections

#### *Salmonella* infection of Caco-2 cells

*Salmonella* and *Legionella* infections were carried out as previously described^25,54,55^. Caco-2 cells (HTB-27; American Type Culture Collection) were maintained in Dulbecco’s modified Eagle’s medium (DMEM) supplemented with 10% (vol/vol) heat-inactivated fetal bovine serum (FBS), 100 IU/mL penicillin and 100 μg/mL streptomycin. Cells were grown at 37°C in a humidified incubator with 5% CO_2_. One day prior to infection, Caco-2 cells were incubated with 0.25% trypsin-EDTA (Gibco) diluted 1:1 with 1 x PBS at 37°C for 15 min to dissociate cells. Trypsin was neutralized with serum-containing medium. 24 hours before infection, cells were replated in DMEM supplemented with 10% (vol/vol) heat-inactivated FBS without antibiotics in a 24-well plate at a density of 3 × 10^5^ cells/well. 3 hours before infection, the media was replaced with Opti-MEM I reduced serum medium (Thermo Fisher Scientific) containing 100 ng/ml Pam3CSK4 (Invivogen) to prime the cells. An overnight culture of wild-type (WT) *Salmonella* Typhimurium (SL1344) was diluted into LB broth containing 300 mM NaCl and then grown for 3 hours at 37°C to induce SPI-1 expression. After induction, the culture was pelleted at 6,010 × g for 3 minutes, washed once with PBS, and then resuspended in PBS. Caco-2 cells were infected with WT *Salmonella* at a multiplicity of infection (MOI) of 60. Control cells were mock-infected with PBS. Cells were centrifuged at 290 × g for 10 min and incubated at 37°C. 1-hour post-infection, infected cells were treated with 100 ng/mL Gentamicin to kill any extracellular *Salmonella* and placed back to 37°C. The supernatants were collected, and cells were lysed by sonication 6 h post infection. Both supernatants and lysates were precipitated by chloroform/methanol and analyzed by immunoblotting.

#### *Legionella* infection of THP1 cells

THP-1 cells in RPMI medium were plated in12-well plates at a density of 8 × 10^5^ cells/well and differentiates with 40 ng/ml phorbol 12-myristate 12-acetate (PMA) for 24 h. The media was replaced with fresh RPMI/10% FBS (without PMA) and the cells were allowed to grow for another 24 h. The media was then replaced with Opti-MEM immediately before *Legionella* infection. The THP-1 cells were infected with a flagellin mutant, Δ*flaA*, of *Legionella pneumophila*^56^, which is an Lp02 strain (*rpsL, hsdR, thyA*) derived from the serogroup 1 clinical isolate Philadelphia-1. *Legionella* was grown as a stationary patch on charcoal yeast extract agar plates at 37°C. After 48 hours, the bacteria were resuspended in PBS and added to the THP-1 cells at a MOI of 10. Infected THP-1s were centrifuged at 400 × g for 10 min and incubated at 37°C for 7 hours. Control cells were mock-infected with PBS. The supernatants and cells were precipitated together by chloroform/methanol and analyzed by immunoblotting.

## Data analysis and statistics

Statistical analysis was performed using GraphPad Prism 9.0 software and Microsoft Excel. Statistical significance was determined using two-sided Student’s *t*-tests.

## Author Contributions

CYT conceived and directed the project, performed experiments, analyzed data, and wrote the manuscript. PE, CH, BD performed experiments and analyzed data. MSE, and JZ performed bacterial infections in Caco-2 cells. JL and TS performed cloning and generated plasmids. All authors read and approved the manuscript.

## Acknowledgements

We thank all the rotation students, Brianna Hill-Payne, Reyna Garcia, and Ivanna Molina for their contributions to the lab set up and discussions. We thank Dr. Igor Brodsky for careful reading of this manuscript and providing feedback. This work was supported by a UNCF/BMS EE Just Early Career Investigator Award and NIH R00 Career Transition Award Grant# 4R00AI148598-03 (CYT) and R01AI123243 and a Burroughs Wellcome Fund Investigator in the Pathogenesis of Infectious Disease Award (SS). MSE is supported by the National Science Foundation Graduate Fellowship DGE-1321851.

## SUPPLEMENTAL FIGURES

**Supplemental Figure S1**.

**Figure S1.**
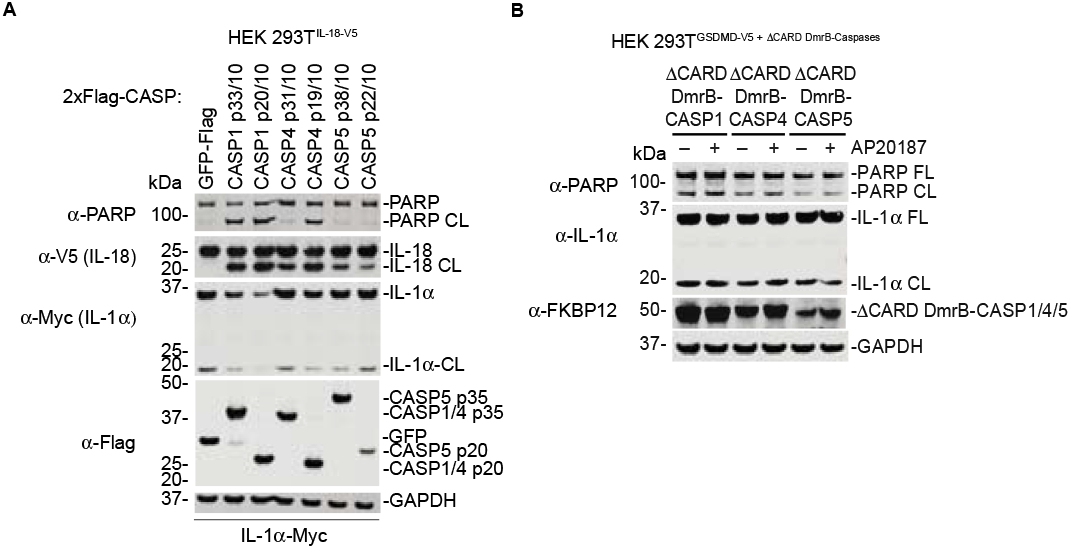
CASP1/4/5 do not cleave IL-1α. (**A**) HEK 293T cells stably expressing IL-18-V5 were transiently transfected with the indicated constructs. After 24 h, samples were then harvested and analyzed by immunoblotting. (**B**) HEK 293T cells stably expressing GSDMD-V5 and either ΔCARD DmrB-CASP1, −4, or −5 were transiently transfected with a plasmid coding for wildtype IL-1α for 24 h. Samples were then treated with AP20187 (1 μM) for 24 h and analyzed by immunoblotting. Data are representative of three or more independent experiments.

